# Ferroptosis contributes to developmental cell death in rice blast

**DOI:** 10.1101/850560

**Authors:** Qing Shen, Meiling Liang, Fan Yang, Yi Zhen Deng, Naweed I. Naqvi

**Author notes:** These authors contributed equally to this work (Qing Shen, Meiling Liang). Author for correspondence: Naweed Naqvi or Yi Zhen Deng.

## Abstract

We identified that ferroptosis, an iron-dependent non-apoptotic cell death process, occurs in the rice blast fungus *Magnaporthe oryzae*, and plays a key role in infection-related development therein. Ferroptosis in the blast fungus was confirmed based on the four basic criteria. We confirmed the dependence of ferroptosis on ferric ions, and optimized C11-BODIPY^581/591^ as a key sensor for subcellular detection and quantification of lipid peroxides that mediate ferroptotic cell death during the pathogenic growth phase of *M. oryzae.* In addition, we uncovered an important regulatory function for reduced glutathione and the NADPH oxidases in generating/modulating the superoxide moieties for ferroptotic cell death in *Magnaporthe*. Ferroptosis was found to be necessary for the specific developmental cell death in conidia during appressorium maturation in rice blast. Such ferroptotic cell death initiated first in the terminal cell and progressed sequentially to the entire conidium. Chelation of iron or chemical inhibition of ferroptosis caused conidial cells to remain viable and led to strong defects in host invasion by *M. oryzae.* Precocious induction of ferroptosis in a blast-susceptible rice cultivar led to resistance against *M. oryzae* invasion. Interestingly, ferroptosis and autophagy were found to play inter-reliant or codependent roles in contributing to such precise cell death in *M. oryzae* conidia during pathogenic differentiation. Our study provides significant molecular insights into understanding the role of developmental cell death and iron homeostasis in infection-associated morphogenesis and in fungus-plant interaction in the blast pathosystem.

## Introduction

Ferroptosis is an iron-dependent form of regulated cell death characterized by membrane damage caused by the accumulation of lipid peroxides to a lethal level (Dixon *et al*., 2012; Dixon & Stockwell, 2014; Kagan *et al*., 2016; Yang *et al*., 2016; Yang & Stockwell, 2016; Stockwell *et al*., 2017; Bersuker *et al*., 2019; Doll *et al*., 2019). Such cell death can be specifically prevented by small molecules such as liproxstatin-1 (Lip-1) and ferrostatin-1 (Fer-1), which were identified in chemico-genetic screens for ferroptosis inhibitors and shown to work as lipophilic antioxidants (Dixon *et al*., 2012; Friedmann Angeli *et al*., 2014). On the other hand, ferroptosis can be induced by glutathione (GSH) depleting reagents such as erastin and buthionine sulfoximine (BSO) through inactivation of glutathione-dependent peroxidases (Yagoda *et al*., 2007; Dixon *et al*., 2012; Yang *et al*., 2014). Using GSH as an essential cofactor, glutathione-dependent peroxidases, particularly glutathione peroxidase 4 (GPX4), reduce peroxidation of phospholipid membranes (Ursini *et al*., 1982; Brigelius-Flohé & Maiorino, 2013; Yang *et al*., 2014). Therefore, GSH depletion with erastin or BSO, or direct GPX4 inhibition with (1S,3R)-RSL3 (hereafter referred to as RSL3), leads to overwhelming lipid peroxidation and causes ferroptotic cell death (Yang *et al*., 2014; Yang *et al*., 2016). Since ferroptosis is a form of iron-dependent oxidative death, iron catalyzed fenton reaction, which generates free radicals, was once speculated as the underlying mechanism (Dixon & Stockwell, 2014; Yang & Stockwell, 2016). However, increasing evidences indicate that iron-dependent enzymes, such as lipoxygenases and NADPH oxidases (NOXs), are the major sources of lipid peroxides that drive ferroptosis (Dixon *et al*., 2012; Kagan *et al*., 2016; Yang *et al*., 2016; Chu *et al*., 2019).

The rice-blast fungus *Magnaporthe oryzae* is a major biotic threat constraining rice production worldwide (Talbot, 2003; Liu *et al*., 2014; Fernandez & Orth, 2018). Given its destructive impact on rice production and well-established molecular genetics, rice blast is the top model system for the investigation of host-pathogen interaction (Talbot, 2003; Ebbole, 2007; Liu *et al*., 2014). The disease starts when the three-celled spore of *M. oryzae*, termed conidium, germinates and produces a polarized germ tube from one of two terminal conidial cells (Fernandez & Orth, 2018). A specialized infection cell, called the appressorium, then forms at the tip of the germ tube. The mature appressorium with a thick melanin layer then uses its accumulated turgor to breach the rice cuticle via a rigid penetration peg and then further proliferates within the rice cells by forming fungal hyphae including penetration and invasive hyphae to establish and spread the disease (Fernandez & Orth, 2018). During appressorium maturation, the three conidial cells and the germ tube transport the storage products within them to the developing appressorium and then collapse and undergo death, although the mechanism driving the conidial death is still not entirely clear (Veneault-Fourrey *et al*., 2006). Autophagy has been reported to be involved in the conidial death, since loss of the essential Atg8 function arrests such cell death and renders *M. oryzae* nonpathogenic, which highlights the close association between conidial cell death and fungal pathogenicity (Veneault-Fourrey *et al*., 2006). In tumor cells, autophagy is found to fine-tune sensitivity to ferroptosis by degradation of the ferritin complex responsible for intracellular iron storage, thereby modulating the content of available iron and the corresponding iron-dependent lipid peroxidation (Bauckman & Mysorekar, 2016; Gao *et al*., 2016; Hou *et al*., 2016; Napoletano *et al*., 2019; Sui *et al*., 2019). However, it remains unknown whether ferroptosis is also involved in the developmental cell death in fungi. *M. oryzae* has a GPX domain containing protein Hyr1, which is required for its full pathogenicity (Huang *et al*., 2011). However, Hyr1, like its yeast orthologue Gpx3, is involved in detoxifying peroxides produced by rice during *in planta* growth (Huang *et al*., 2011). Therefore, whether *M. oryzae* also has a GPX4-like reducing enzyme involved in internal lipid peroxidation is still not known. What is known is that *M. oryzae* has lipid peroxide producing NADPH oxidases, Nox1, Nox2 and Nox3, with the former two being highly similar and better characterized (Egan *et al*., 2007; Ryder *et al*., 2013). Nox2 and Nox1 are required for pathogenesis by organizing the penetration peg formation and elongation of penetration hyphae during rice infection, respectively (Ryder *et al*., 2013). However, both genes, in particular *NOX2*, have relatively high expression during appressorium maturation (Ryder *et al*., 2013), suggesting their additional roles before rice penetration. Given the contribution of mammalian NOX enzymes to ferroptosis-inducing lipid peroxidation, it is interesting to investigate whether the Nox proteins in *M. oryzae* also control the cell death in conidia through ferroptosis mechanism.

*M. oryzae* is a typical hemibiotrophic fungus that first establishes an intimate biotrophic association within the first invaded epidermal cell in the susceptible host and subsequently develops infection hyphae to spread into and kill the neighbouring cells/tissue after it switches to necrotrophic growth (Fernandez & Orth, 2018). At both biotrophic and necrotrophic stages, *M. oryzae* copes with strong host defenses such as the production of reactive oxygen species (ROS) including peroxides, which can strengthen rice cell walls as well as kill the invading fungus directly via highly coordinated programmed cell death (Bradley *et al*., 1992; Levine *et al*., 1994; Chi *et al*., 2009; Parker *et al*., 2009; Huang *et al*., 2011; Wang *et al*., 2019). When *M. oryzae* interacts with a blast-resistant rice cultivar (termed incompatible interaction), the host immune response, hypersensitive response (HR) cell death in particular, is extremely strong, and blocks *M. oryzae* from invading the host and thus stops the establishment of blast disease (Liu *et al*., 2014). In a seminal study, ferroptotic cell death was reported to be involved in the HR response in the presence of ROS-generating NADP-malic enzyme 2 during incompatible rice-*M. oryzae* interaction (Dangol *et al*., 2019), although further experiments specifically focusing on the rice side are required to validate these findings. Considering the general requirement of ROS production to rice immunity and the universal occurrence of ferroptosis in diverse biological contexts (Distéfano *et al*., 2017; Stockwell *et al*., 2017; Conrad *et al*., 2018; Dangol *et al*., 2019), it is interesting to test whether ferroptosis is generally involved in rice defense against *M. oryzae* infection, in particular whether inducing ferroptosis in susceptible rice can switch on its resistance against the invading pathogen.

In this study, we report that ferroptosis contributes to conidial cell death during the infection-related morphogenesis in *M. oryzae*. We also found that the ferroptotic cell death occurs sequentially and is first initiated in the terminal cell, and later progresses to the other two conidial cells during appressorium formation and maturation, and such ferroptotic cell death is required for *M. oryzae* infection in rice. On the other hand, we also found that induction of ferroptosis in a susceptible rice cultivar leads to blast resistance against *M. oryzae* invasion. Overall, our findings suggest that ferroptosis is a conserved cell death mechanism employed by both partners involved in the rice blast pathosystem for developmental and immune purposes, respectively. We provide evidence that ferroptosis, in concert with ROS, not only contributes to rice resistance to *M. oryzae* infection but also positively regulates *M. oryzae* pathogenic development and pathogenicity. Lastly, we show that ferroptosis intersects with as well as acts in parallel with autophagy for proper induction of conidial cell death. Our study contributes significantly to understanding the role of iron homeostasis in fungus-plant interaction, and will be helpful in developing strategies for management and control of rice blast using ferroptosis pathway and/or the superoxide moieties as crucial molecular targets for disease control.

## Materials and Methods

### Fungal strains and growth conditions

*M. oryzae* wild-type strain B157 was obtained from the Directorate of Rice Research (Hyderabad, India), and was used as the parental strain for all the experiments and transformations in this study. The *H1-mCherry*, *atg8*Δ and the complemented *atg8*Δ strains have been described in our previous reports (Deng *et al*., 2009; Yadav *et al*., 2019), while the *atg8*Δ strain complemented with *GFP-ATG8* was kindly provided by Dr. Kou Yanjun. Other strains were generated through *Agrobacterium tumefaciens*-mediated transformation and confirmed by locus-specific PCR as described (Rho *et al*., 2001; Ramanujam & Naqvi, 2010; Yang & Naqvi, 2014). The corresponding plasmid vectors were constructed as follows: pHistone H1-GFP was constructed by inserting PCR amplified fragments of histone H1 ORF (without the stop codon), GFP open reading frame (ORF) without the start codon, and 3’UTR sequence of histone H1 sequentially into the backbone vector pFGL822 (Addgene #58225, Basta resistance); pCytoplasm GFP was constructed by inserting PCR fragments of histone H3 promoter and the GFP ORF sequentially into the backbone vector pFGL1010 (Addgene #119081, sulfonylurea resistance); *NOX1* and *NOX2* deletion constructs were generated by inserting PCR amplified 5’ UTR and 3’ UTR fragments of the target genes sequentially into backbone vector pFGL821 (Addgene #58223, hygromycin resistance), flanking the *HPH* (*hygromycin B phosphotransferase*) gene. The primers used for plasmid construction are summarized in Table S1. Except for transformants, all the *M. oryzae* strains were propagated on Prune-agar medium as described (Ramanujam & Naqvi, 2010).

### Chemical treatment and cell viability measurement

Freshly harvested conidia at a concentration of 1 × 10^5^ conidia/ml in sterile water were inoculated on hydrophobic cover glass (Menzel-Glaser), with or without treatment of the following chemicals individually: 5 *µ*M FeCl_3_, 100 *µ*M BSO (Sigma, B2515), 5 *µ*M FeSO_4_, 5 *µ*M CuSO_4_, 10 *µ*M erastin (Sigma, E7781), or 4 *µ*M RSL3 (Sigma, SML2234). The abovementioned chemicals were added at 0 hours post inoculation (hpi), and cell viability was quantified at 14.5 hpi. For the conidia treated with 5 *µ*M ciclopirox olamine (Sigma, C0415), 54 *µ*M Lip-1 (Sigma, SML1414), 25 *µ*M diphenyleneiodonium (Sigma, D2926), 5 mM deferoxamine (Sigma, D9533), 5 *µ*M Fer-1 (Sigma, SML0583), or 5 mM GSH (Sigma, G4251), the chemicals were added at 4 hpi, and cell viability was quantified at 24 hpi. For co-treatments, FeCl_3_ or BSO was added at 0 hpi, and 4 hours later Lip-1 or CPX was added. Unless indicated otherwise, the methodology for conidial inoculation and chemical treatment, in particular chemical concentration and time points to add the chemicals, was always the same. The cell viability was evaluated either by examining epifluorescence of intact nuclei (visualized by hH1-GFP or hH1-mCherry), or by staining with 1% trypan blue (Sigma, T6146) solution at room temperature for 20 min before microscopic examination.

### Staining procedures and microscopy

Conidia on cover slips were inoculated with 10 *µ*M C11-BODIPY™581/591 (Thermo Fisher, D3861) for detection of lipid peroxides, 1 *µ*M calcein-AM (Invitrogen, C3099) for iron staining, 2 *µ*g/ml fluorescein diacetate (Sigma, F7378) for viable cell staining, or 28 *µ*M Cell Tracker™ Blue CMAC (Invitrogen, C2110) for vacuolar staining at room temperature for 20-30 min, followed by two washes with sterile water.

Epifluorescence was observed using a Leica TCS SP8 X inverted microscope system equipped with an HCX Plan Apochromat 63×/1.2 water objective (Leica Microsystems). Excitation of fluorescent dye or fluorescently labelled proteins was carried out using the argon laser (488nm/65mW) for C11-BODIPY and GFP-Atg8 (emission, 500-550 nm); the white light laser for calcein-AM (excitation, 488 nm; emission, 510-560 nm), FDA (excitation, 488 nm; emission, 500-550 nm), and H1-mCherry (excitation, 543 nm; emission, 610-680 nm); and the diode laser for CMAC (excitation, 405 nm; emission, 410-480 nm). All the lasers were controlled by 8 channel AOTF (Acousto-Optical-Tunable-filter) for rapid modulation of laser intensity. Images were captured using the Hybrid Detector of this system. All parts of the system were under the control of its own Leica Application Suite X software package. Epifluorescence of H1-GFP or the cytoplasmic GFP was observed using a LSM5 Exciter microscope system equipped with an argon laser (excitation, 488 nm; emission, 505-550 nm) (Carl Zeiss). GFP-Atg8 subcellular localization was shown as 3D-rendered images, whereas others were shown as single layer images or maximum intensity projections of five optical slices as indicated.

### MDA measurement

Conidia inoculated on cover glass (Menzel-Glaser, 80×80 mm) were treated with solvent, or Lip-1, or FeCl_3_, or ciclopirox olamine, or diphenyleneiodonium and harvested at 14.5 hpi. Two hundred million conidia were used for each treatment. Samples were snap-freezed and ground to a fine powder using liquid nitrogen with mortar and pestle. MDA levels were measured using a TBARS (TCA Method) Assay Kit (Cayman, 700870) following the manufacturer’s instructions.

### Blast infection assays

Rice seedlings (var. CO39) grown in the greenhouse up to the four-leaf stage were used for blast infection assays. Rice leaf sheath infection was performed as described (Deng *et al*., 2012; Selvaraj *et al*., 2017). For sheath infections focusing on *M. oryzae* pathogenicity, the way of doing chemical treatment was the same as described before, except that all the chemicals were washed off and replaced by sterile water at 22 hpi. To induce ferroptosis in rice, RSL3 or erastin were applied at 22 hpi, and rice resistance was examined at 48 hpi. DAB (3,3’-diaminobenzidine; Sigma, D8001) staining was performed as described (Sun *et al*., 2006; Daudi & O’Brien, 2012).

### Iron measurement

*M. oryzae* conidia were harvested and normalized using the Iron Assay Buffer (100 ml) from the Iron Assay kit (Abcam, ab83366), to reach a final conidia count of 3×10^5^/well. Setting up of standard curve, inoculation with Iron Probe and/or Iron Reducer, measurement of Fe^2+^ or total iron (Fe^2+^ + Fe^3+^) were performed following the manufacturer’s instructions, and using a colorimetric microplate reader (Biotek, Synergy H1; OD=593 nm).

## Results

### Ferroptosis contributes to developmental cell death in *M. oryzae*

To investigate the involvement of ferroptosis in the conidial death observed in *M. oryzae*, we first tested whether the precise cell death is iron dependent by treating the conidia of a nucleus marked wild-type strain (*H1-GFP*) with the membrane-permeable iron chelator ciclopirox olamine (CPX). The conidial death, indicated by breakdown of nuclei, collapse of the conidial cells, and accumulation of the vital dye trypan blue, was dramatically suppressed at 24 hpi by CPX (Fig. 1a-c; *P* < 0.01, n = 900 conidia). Similar cell death suppression effect on the conidia, although not as robust as CPX, was also evident when another iron chelator deferoxamine (DFO) was used (Fig. S1a,b; *P* < 0.01, n = 900 conidia). To confirm the involvement of ferroptosis in the conidial death, the wild-type conidia were further treated with a panel of established ferroptosis inhibitors or inducers (Dixon *et al*., 2012; Friedmann Angeli *et al*., 2014; Yang *et al*., 2014; Stockwell *et al*., 2017). The conidial death was suppressed by the two specific inhibitors of ferroptosis, Lip-1 or Fer-1 (Figs 1c,d, S1c,d; *P* < 0.01, n = 900 conidia), with Lip-1 clearly showing a dose-dependent suppression efficiency at 24 hpi (Fig. 1d; *P* < 0.01, n = 900 conidia). Addition of iron (FeCl_3_) or the GSH biosynthesis inhibitor BSO, which has been reported to induce ferroptosis (Dixon *et al*., 2012; Yang *et al*., 2014), advanced the conidial death at 14.5 hpi (Figs 1e-g, S1f,g; *P* < 0.05, n = 900 conidia). Exogenous iron-promoted sensitivity to the conidial death was reverted upon co-treatment with Lip-1, and attenuated by co-treatment with the iron chelator CPX (Fig. 1e; *P* < 0.05, n = 900 conidia). Similarly, CPX arrested the conidial cell death, regardless of the BSO treatment (Fig. 1g; *P* < 0.01, n = 900 conidia). Direct addition of GSH suppressed the conidial death (Fig. S1e; *P* < 0.01, n = 900 conidia) but surprisingly promoted death in the appressoria (Fig. S1e), thus suggesting a precise control of GSH/ROS levels and the sites of action during *M. oryzae* development. Together, the conidial death in response to ferroptosis inhibitors or inducers indicates that the iron-dependent cell death in the conidium is ferroptotic in nature.

**Fig. 1.**
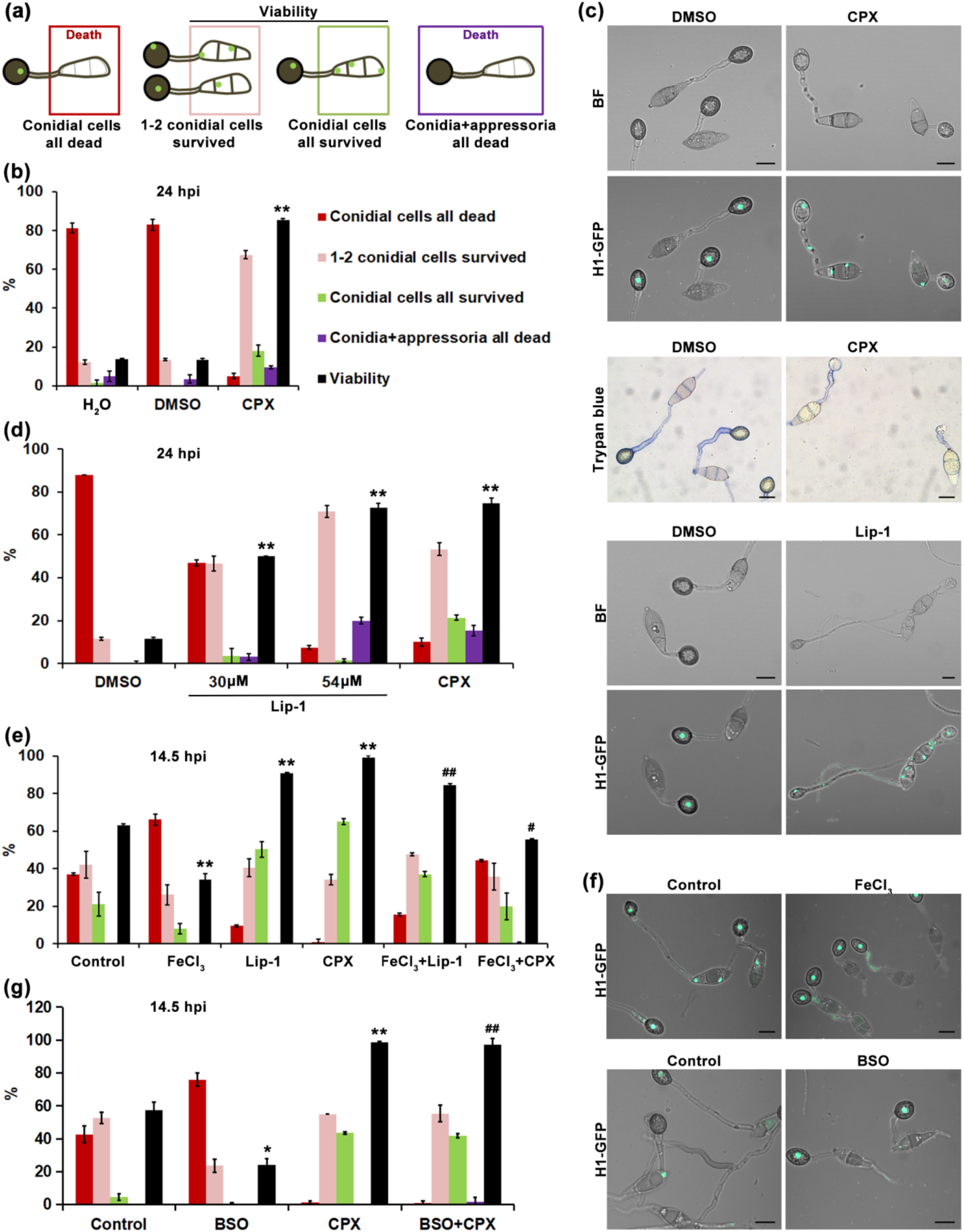
Ferroptosis contributes to conidial death in *M. oryzae*. (a) Schematic representation of the conidial cells all dead (red), 1-2 conidial cells survived (pink), conidial cells all survived (green) and conidia + appressoria all dead (purple). The Viability (black) is the sum of pink and green (the 1-2 conidial cells survived and conidial cells all survived). Cell viability was assessed based on whether an intact nucleus (visualized by a nucleus-marked *HistoneH1-GFP* strain) was present, as quantified in Fig. 1b, d, e, g. (b) Ability of iron chelator CPX to prevent conidial death. Conidial viability was measured 24 hpi. Mean ± s.d. from three independent replicates (one hundred conidia per replicate) is shown. ** *P* < 0.01versus DMSO control (*t*-test). (c) Visualization of conidial viability in the presence of CPX or Lip-1 in *H1-GFP* strain at 24 hpi using epifluorescence microscopy (upper and bottom panel) or trypan blue staining (middle panel). BF, bright field. H1-GFP signal represents the maximum intensity projection for five optical slices. Bar, 10 *µ*m. (d) Ability of Lip-1 or CPX to prevent conidial death. Conidial viability was measured at 24 hpi using the *H1-GFP* strain. Mean ± s.d. from three independent replicates (100 conidia per replicate) is shown; ** *P* < 0.01 versus DMSO control (*t*-test). (e) Ability of FeCl_3_ to promote conidial death, and the suppression effect of Lip-1 or CPX on conidial death promoted by FeCl_3_. Conidial viability was measured at 14.5 hpi, and are plotted as the mean ± s.d. (3 replicates, 100 conidia for each replicate); ** *P* < 0.01 versus control, # *P* < 0.05 and ## *P* < 0.01 versus FeCl_3_ (*t*-test). (f) Visualization of conidial viability in the presence of FeCl_3_ or BSO in the *H1-GFP* strain at 24 hpi using epifluorescence microscopy. Scale bar represents 10 **µ**m. (g) Ability of BSO to promote conidial death, and the suppression effect of CPX on conidial death promoted by BSO. Conidial viability was measured at 14.5 hpi using the *H1-GFP* strain. Mean ± s.d. from three independent replicates (100 conidia each replicate) is shown; * *P* < 0.05 and ** *P* < 0.01 versus control, ## *P* < 0.01 versus BSO (*t*-test). All data are representative of three independent experiments (biological replicates). Total n=900 conidia were used to generate the data interpretation as indicated in the main text.

Next, we sought to validate whether such iron-dependent conidial death is associated with an increase in lipid peroxides, a defining event and a hallmark of ferroptosis. Lipid peroxides, as detected by the lipid peroxidation fluorescent probe C11-BODIPY^581/591^ (Dixon *et al*., 2012; Friedmann Angeli *et al*., 2014; Yang *et al*., 2014), accumulated intensely in the germinating terminal cell and weakly in the other two cells of the conidium when observed at 7 hpi (Fig. 2a), which was deemed as the initiation time point for the conidial death to ensue (Fig. 3). Exogenous addition of BSO or iron (FeCl_3_) showed an increase in C11-BODIPY fluorescence in the terminal conidial cells and helped to spread the lipid peroxidation to the remaining two cells (Fig. 2a). In contrast, both CPX and Lip-1 caused a large drop in C11-BODIPY fluorescence, especially in the conidial cells (Fig. 2a). The membrane associated accumulation of C11-BODIPY in the conidium and the consequent change of fluorescence in response to ferroptosis induction or suppression strongly implicate the detected lipid peroxides as inducers of ferroptosis. Whereas, the cytosolic C11-BODIPY signal in the germ tubes and maturing appressoria likely results from lipid peroxidation events required for proper development, and did not show significant response to the ferroptosis inhibitor Lip-1 (Fig. 2a). Loss of C11-BODIPY fluorescence within the mature appressoria was frequently observed (Figs 2a, 3a, 4f, 6b, S3b) and is likely due to masking by the melanin layer, since similar loss of fluorescence was also noticed in mature appressoria of the wild-type strain when a vital dye calcein-AM was applied (Fig. S3a). We further quantified the endogenous lipid oxidation during the iron-dependent conidial death by measuring the accumulation of malondialdehyde (MDA), a common product from the oxidative degradation of multiple lipid species (Gaschler & Stockwell, 2017; Gaschler *et al*., 2018). Lip-1 or CPX treated wild-type conidia showed a decrease in MDA accumulation, whereas treatment with iron (FeCl_3_) showed the opposite effect and an increase in MDA levels at 14.5 hpi (Fig. 2b; *P* < 0.01, n ≥ 0.6 billion conidia). Together, the C11-BODIPY oxidation and the MDA quantification data indicate an iron-dependent lipid peroxidation event occurring in the conidia undergoing cell death. However, erastin and RSL3, the two well-known inducers of ferroptosis in mammalian cells, did not affect conidial cell death or lipid peroxidation levels as probed by C11-BODIPY (Figs 6a,b, S1i; *P* > 0.05, n = 900 conidia). Erastin causes GSH depletion by inhibiting the cysteine/glutamate antiporter system x_c_^-^, while RSL3 inhibits GPX4 through direct binding. Protein BLAST found no orthologs of system x_c_^-^ components in *M. oryzae.* Thus, the plant ROS-targeting Hyr1 remains the only *M. oryzae* locus related to GPX4, which likely explains why these two chemicals were unable to induce ferroptosis in *M. oryzae*. Collectively, these findings led us to conclude that ferroptosis occurs in *M. oryzae* conidia, and contributes to the developmental cell death therein.

**Fig. 2.**
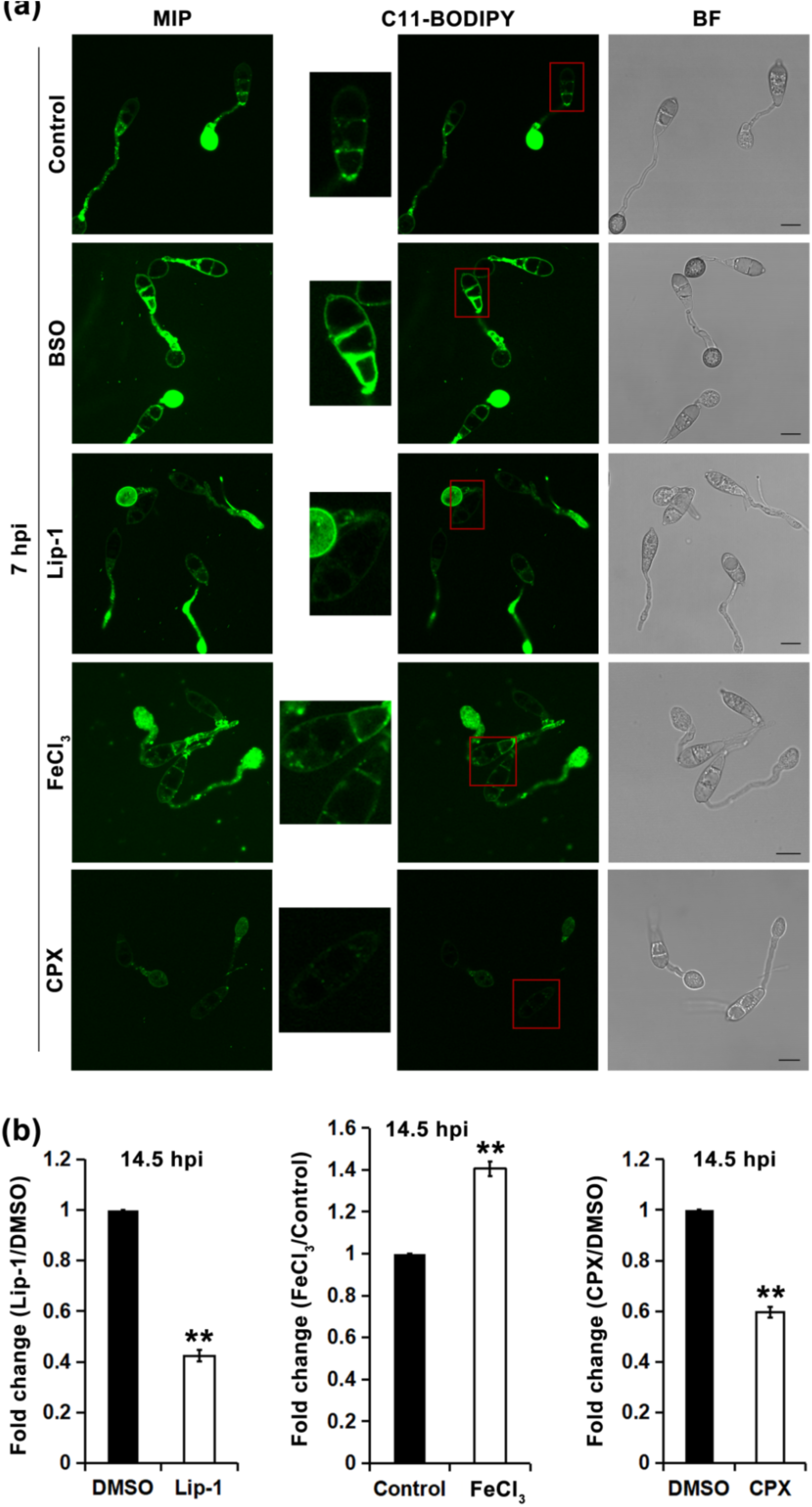
Increase of lipid peroxidation marks the occurrence of ferroptosis in the conidium. (a) Ability of conidium-death-modulating compounds (BSO, Lip-1, FeCl_3_, and CPX) to regulate C11-BODIPY581/591 oxidation in wild-type *M. oryzae* conidia. Epifluorescent microscopy was performed using conidia inoculated on inductive surface (cover slip) for 7 h, followed by staining with C11-BODIPY581/591. Red rectangles mark the sites of origin for the respective insets. MIP, maximum intensity projection. C11-BODIPY, oxidized C11-BODIPY581/591. BF, bright field. Representative images shown for each treatment were from three independent replicates with similar results. Bar = 10 **µ**m. (b) Ability of Lip-1, FeCl_3_, and CPX to modulate MDA accumulation in wild-type *M. oryzae*. MDA levels were measured at 14.5 hpi and are presented as fold change of chemical treated versus mock control. Data are mean ± s.d. from three to four technical replicates (two hundred million conidia per treatment per replicate); ** *P* < 0.01 versus DMSO/solvent control (*t*-test). Representative of three independent experiments (biological replicates) is shown, and total n ≥ 0.6 billion conidia were used to generate the data interpretation as indicated.

**Fig. 3.**
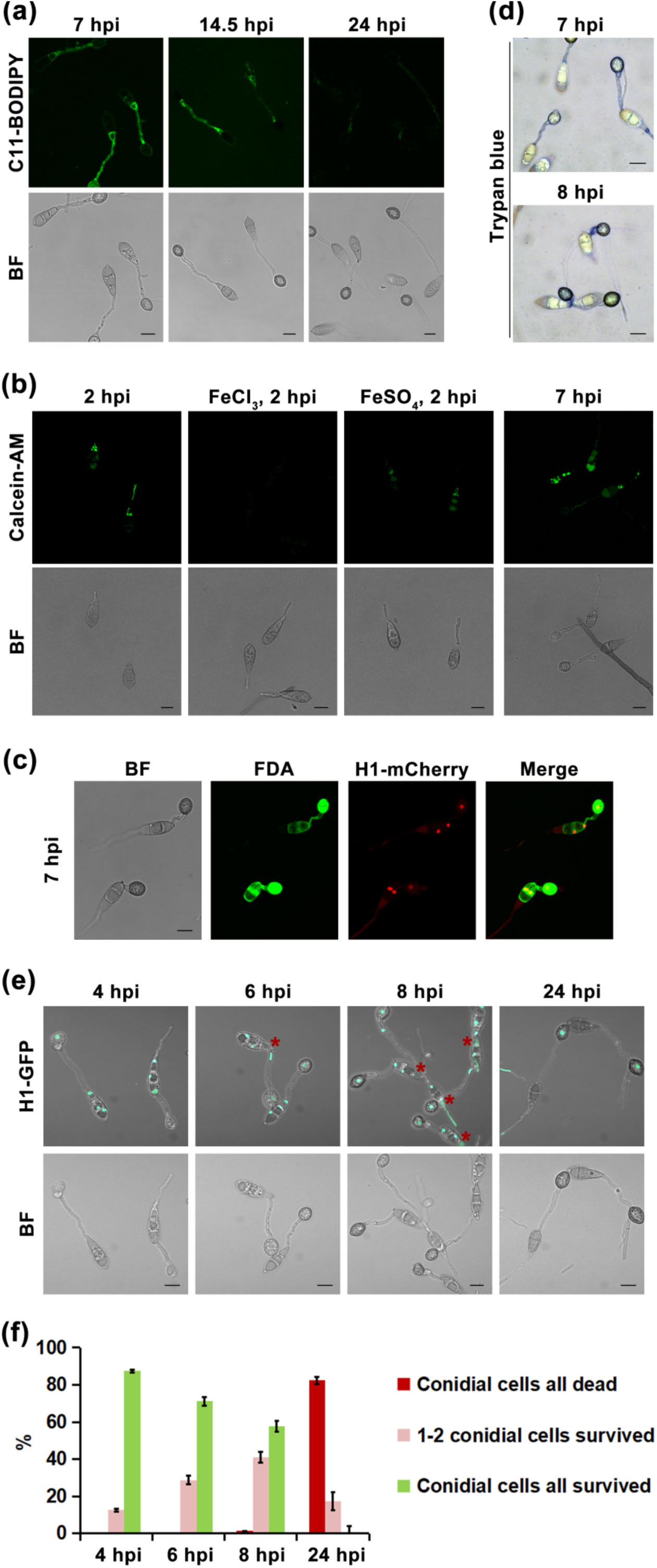
Terminal conidial cell initiates ferroptosis first. (a) Lipid oxidation visualized by C11-BODIPY581/591 staining in wild-type *M. oryzae* conidium at 7 hpi, 14.5hpi, and 24 hpi. BF, bright field. (b) Cellular iron levels in *M. oryzae*. Wild-type conidial cells cultured in the presence or absence of FeCl_3_ (25**µ**M) or FeSO_4_ (25**µ**M) were probed with calcein-AM, and fluorescence observed at 2 or 7 hpi. BF, bright field.(c) FDA staining of the *HistoneH1-mCherry* strain showing death of terminal conidial cells at 7 hpi. Merge, merge of FDA and H1-mCherry. (d) Death of terminal conidial cells in wild-type *M. oryzae* shown by trypan blue staining (e) Visualization of *H1-GFP* conidial death at the indicated time points using epifluorescence microscopy. BF, bright field. Red asterisks mark the dead terminal conidial cells. (f) Bar chart quantifying the sequential cell death in conidial cells. Mean ± s.d. is shown (3 replicates, 100 conidia per replicate). Except for BF and trypan blue, all images are shown as maximum intensity projections. Scale bars equal 10 **µ**m in each instance. All data are representative of two to three independent experiments (biological replicates).

**Fig. 4.**
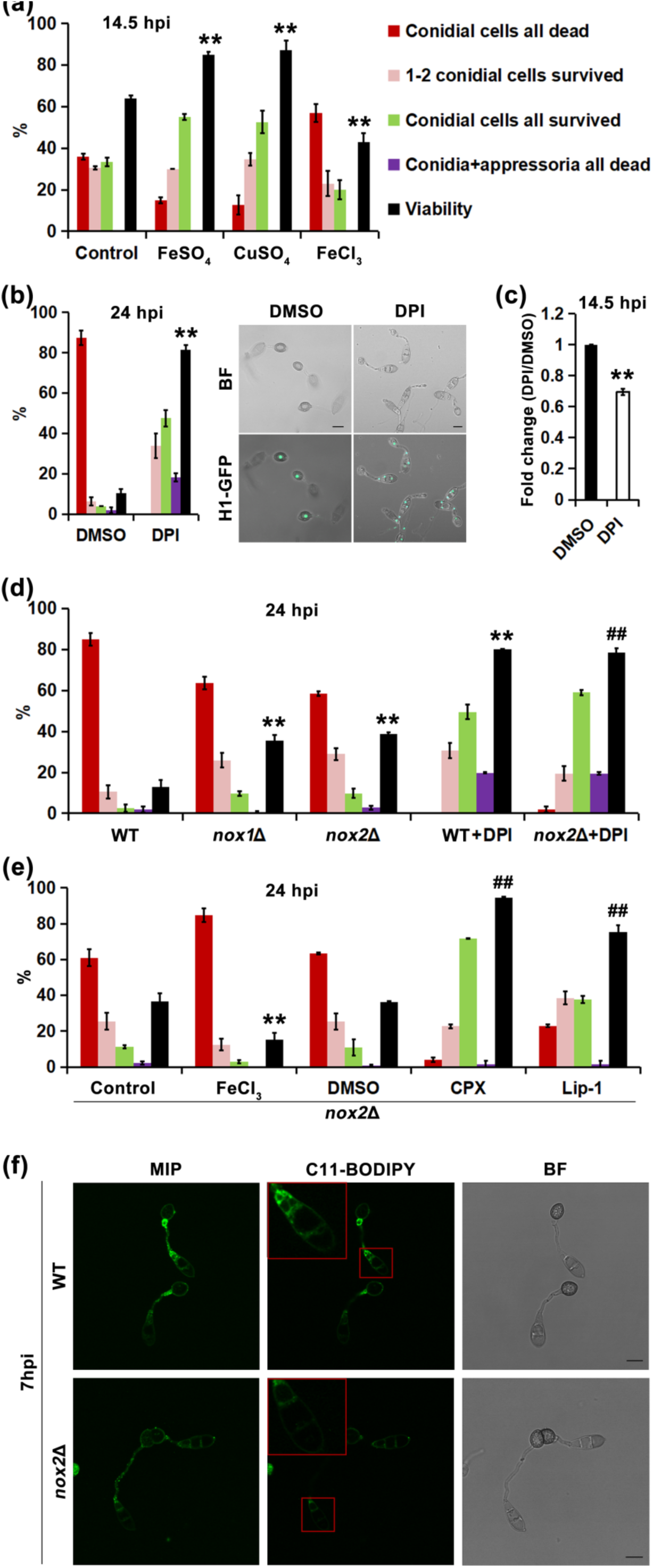
Nox enzymes are a source of ferroptosis-inducing lipid peroxidation in *M. oryzae*. (a) Conidial viability in the *HistoneH1-GFP* conidial cells treated with FeSO_4_, CuSO_4_, or FeCl_3_. Conidial viability was quantified at 14.5 hpi, and is plotted as the mean ± s.d. (100 conidia per replicate, 3 independent replicates,); ** *P* < 0.01 versus control (*t*-test). (b) Quantification (left panel) and visualization (right panel) of conidial death prevented by DPI treatment at 24 hpi using the *H1-GFP* strain. Mean ± s.d. from three independent replicates (100 conidia for each) is shown. ** *P* < 0.01 versus DMSO (*t*-test). BF, bright field. H1-GFP signal is shown as the maximum intensity projections. Bar =10 **µ**m. (c) Ability of DPI to prevent MDA accumulation in wild-type *M. oryzae* at 14.5 hpi. Data are mean ± s.d. (four technical replicates, two hundred million conidia per replicate) presented as fold change of DPI treated versus DMSO; ** *P* < 0.01 versus DMSO (*t*-test). (d) Conidium viability of wild-type and *nox*Δ mutants at 24 hpi, in the presence or absence of DPI. Mean ± s.d. (3 replicates, 100 conidia for each replicate) is shown; ** *P* < 0.01 versus B157, ## *P* < 0.01 versus *nox2*Δ (*t*-test). WT, wild-type. (e) Conidium viability of *nox2*Δ in the presence of FeCl_3_, CPX, or Lip-1 (30 **µ**M). Mean ± s.d. is shown (3 replicates, 100 conidia for one replicate,); ** *P* < 0.01 and ## *P* < 0.01 versus *nox2*Δ control and DMSO, respectively (*t*-test). (f) Lipid oxidation visualized by BODIPY 581/591 C11 staining in conidial cells of wild-type and *nox2*Δ at 7 hpi. Red rectangles mark the origin of the highlighted insets. MIP, maximum intensity projection. C11-BODIPY, oxidized C11-BODIPY581/591. BF, bright field. WT, wild-type. Data represents two to three independent experiments (biological replicates), n=900 or n=600 conidia in total were used for each data interpretation as indicated.

### Sequential conidial cell death during proper appressorium formation and function

As mentioned earlier, lipid peroxides showed higher accumulation in one of the terminal cells prior to spreading to the other two conidial cells (Figs 2a, 3a), suggesting that the three conidial cells do not die simultaneously. Instead, the terminal conidial cell initiates ferroptotic death first. Supporting this hypothesis, less free iron was detected in the terminal conidial cells before (2 hpi) and during (7 hpi) conidial death (Fig. 3b) as judged by using calcien-AM, which is a probe for iron and also serves the dual function of being a vital dye too (Thomas *et al*., 1999; Yang *et al*., 2016). Fluorescence of calcein-AM was quenched dramatically and moderately upon binding to ferric and ferrous ions, respectively (Fig. 3b), thus underscoring its role as a specific probe for iron. Based on calcein-AM, less iron was deemed to be available in the terminal conidial cells probably because they are occupied by lipid peroxide producing enzymes, such as Nox1 and Nox2, for subsequent ferroptosis initiation. To assess whether conidial cells undergo ferroptosis sequentially, we tracked conidial cell death at multiple time points including 4 hpi, 6 hpi, 7 hpi, 8 hpi, and 24 hpi during pathogenic development. Indeed, a small percentage of terminal conidial cells started to die as early as 6 hpi (Fig. 3f; n = 600 conidia). The death of the terminal conidial cell, further supported by loss of fluorescence of the cytoplasmic vital dye fluorescein diacetate (FDA), which detects metabolically active cells (Jones *et al*., 2016), was more obvious and abundant at 7 and 8 hpi (Fig. 3c-f). Overall, these results helped us infer that the conidial cells initiate ferroptosis sequentially with precedence accorded to the terminal cell.

### Nox enzymes are a potential source of ferroptosis-inducing lipid peroxidation in *M. oryzae*

We found that ferric ion, but not ferrous ion or copper which also catalyses the fenton reaction, sensitized the conidia towards cell death (Fig. 4a; *P* < 0.01, n = 900 conidia). Therefore, fenton chemistry is not the major source of ferroptosis-inducing lipid peroxidation in *M. oryzae*. Next, we examined the involvement of membrane-associated NADPH oxidases or Nox, mammalian orthologs of which have been reported in ferroptosis regulation (Dixon *et al*., 2012), in ferroptotic conidial death in rice blast. The conidial ferroptosis was strongly prevented by the canonical Nox inhibitor diphenylene iodonium (DPI) and partially inhibited upon loss of Nox1 or Nox2 function through gene deletion (hereafter *nox2*Δ#5 chosen as representative *nox2*Δ) (Figs 4b,d, S1h; *P* < 0.01, n = 600 conidia). Moreover, conidial viability in *nox2*Δ could be further enhanced upon addition of DPI (Fig. 4d; *P* < 0.01, n = 600 conidia), thus suggesting that Nox1 and Nox2 act redundantly in regulating ferroptotic cell death in *M. oryzae*. Given the relatively higher expression of *NOX2* during conidial death (Ryder *et al*., 2013), *nox2*Δ was chosen for further characterization. The reduction of Nox activity was further verified by a decrease in MDA accumulation with DPI inhibition and a decline in C11-BODIPY fluorescence in *nox2*Δ, respectively (Fig. 4c,f; *P* < 0.01, n = 0.4 billion conidia). Notably, conidial viability in *nox2*Δ was undetectable when iron (FeCl_3_) was applied, whereas CPX and Lip-1 strongly enhanced it (Fig. 4e; *P* < 0.01, n = 600 conidia), which further supports the conclusion that the Nox2-enabled conidial death is indeed ferroptotic. We conclude that *M. oryzae* Nox proteins, like the mammalian counterparts, participate in ferroptosis, and modulate the cell death in conidia during infection-related morphogenesis.

### Ferroptotic fungal cell death is required for pathogenesis

To investigate the importance of ferroptotic conidial death to the pathogenicity of *M. oryzae*, we inoculated the wild-type B157 strain or a cytoplasmic GFP-marked wild-type strain in the presence of ferroptosis inhibitors or inducer on the susceptible rice cultivar CO39. To better define the chemical effect on conidial death, these chemicals were removed at 22 hpi prior to fungal invasion of the rice leaf. Both Lip-1 and CPX, which function as inhibitors of ferroptosis, caused a huge drop in the ability to develop fungal penetration/infection hyphae within the rice cells at 28 hpi (Figs 5a-d, S2a; *P* < 0.01, n ≥ 900 conidia). Further observation at 32 hpi and 48 hpi showed that *M. oryzae*, whose conidial death was suppressed before invasion, still retained the ability to penetrate and form fungal hyphae within the host, but the overall process was greatly delayed (Fig. 5b-d; *P* < 0.01, n = 1200 conidia). Similar decrease in ability to develop fungal hyphae *in planta* was observed in *nox2*Δ, and in wild-type *M. oryzae* treated with DPI before 22 hpi to rule out the role of Nox2 in penetration peg formation (Figs 5e,f, S2; *P* < 0.01, n = 1200 conidia). Conversely, excess iron (FeCl_3_), which functions as an inducer of ferroptosis, advanced the *in planta* hyphal development (Fig. 5a,b; *P* < 0.01, n = 900 conidia). Overall, these findings helped us infer that proper ferroptotic cell death in the conidium is necessary for ensuring robust infection by *M. oryzae*.

**Fig. 5.**
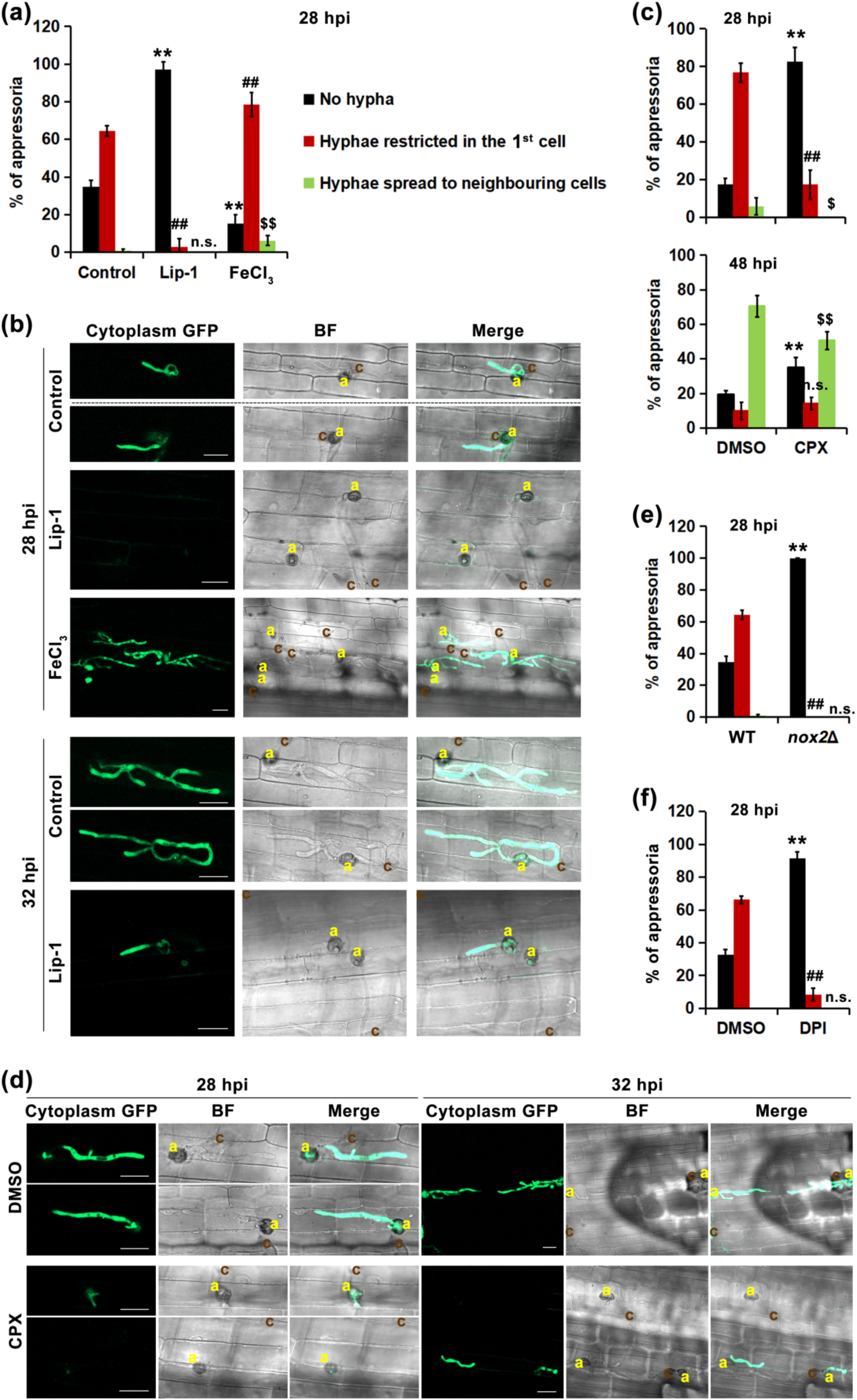
Ferroptotic conidial death is required for *M. oryzae* pathogenicity. (a) Ability of wild-type *M. oryzae* treated with Lip-1 or FeCl_3_ to form fungal hyphae within rice cells at 28 hpi. Data are mean ± s.d. from three independent replicates (100 conidia per replicate); ** *P* < 0.01, ## *P* < 0.01 and $$ *P* < 0.01 versus “no hypha”, “hyphae restricted in the 1^st^ cell”, and “hyphae spread to neighbouring cells” of control, respectively; n.s., not significant (*t*-test). (b) Microscopic observation of hyphal growth within rice cells at 28 and 32 hpi using the cytoplasmic GFP-expressing strain. BF, bright field. Merge, merge of cytoplasm GFP and BF. C, conidia; A, appressorium. (c) Quantification of hypha formation in rice at 28 hpi and 48 hpi using the wild-type *M. oryzae* strain in the presence of CPX. Data are mean ± s.d. from four independent replicates (100 conidia for each replicate); ** *P* < 0.01, ## *P* < 0.01, $ *P* < 0.05 and $$ *P* < 0.01 versus “no hyphae”, “hyphae restricted in the 1^st^ cell”, and “hyphae spread to neighbouring cells” of corresponding DMSO, respectively; n.s., not significant (*t*-test). (d) Ability of the *cytoplasm GFP* strain treated with CPX to develop fungal hyphae within rice cells. Epifluorescent microscopy was performed at 28 and 32 hpi. BF, bright field. Merge, merge of cytoplasm GFP and BF. C, conidia; A, appressorium. (e) Ability of wild-type and *nox2*Δ to form infection hyphae within rice cells at 28 hpi. Mean ± s.d., 4 replicates (100 conidia for each replicate); ** *P* < 0.01 and ## *P* < 0.01 versus “no hyphae” and “hyphae restricted in the 1^st^ cell” of wild-type, respectively; n.s., not significant (*t*-test). WT, wild-type. (f) Effect of DPI inhibition on infection hyphae formation within rice cells at 28 hpi. Mean ± s.d. from 4 replicates (100 conidia per replicate) is shown; ** *P* < 0.01 and ## *P* < 0.01 versus “no hyphae” and “hyphae restricted in the 1^st^ cell” of DMSO, respectively; n.s., not significant (*t*-test). All the chemicals were washed away at 22 hpi, a time-point before invasive growth initiates, so that the effect of the chemicals was specific/restricted to conidial death but not to rice cell death related to HR. Scale bars equal 20 **µ**m in each instance. All data are representative of three independent biological replicates, and n=900 or n=1200 conidia in total were used for the data interpretation as indicated in the main text.

### Inducing ferroptosis in a susceptible rice cultivar leads to blast resistance

Since ROS production and regulated cell death are reported to be involved in host resistance to *M. oryzae* infection (Dangol *et al*., 2019), we were also interested in assessing the consequences of ferroptosis induction in a blast-susceptible rice cultivar. Towards this goal, we applied RSL3 or erastin, the two ferroptosis inducers that do not affect conidial cell death (Figs 6a,b, S1i), at 22 hpi to the blast-sensitive CO39 rice sheath inoculated with the wild-type *M. oryzae* isolate, and examined the fungal hyphal development therein. Without RSL3 treatment, most wild-type hyphae successfully spread to rice cells surrounding the site of initial penetration at 48 hpi (Fig. 6c,d). However, RSL3 application led to a failure in forming hyphae within the blast-susceptible rice sheath, and caused a significant drop in the ability of the infection hyphae to spread to the neighbouring cells (Fig. 6c,d; *P* < 0.01, n = 900 conidia). In addition, the few hyphae that attempted to spread to the next cells elicited a strong immune response from the host in the first invaded cells (Fig. 6d). Similar failure to develop fungal hyphae within the penetrating rice cells and their surrounding cells was observed when erastin was used (Fig. 6c,d; *P* < 0.01, n = 900). In addition, without RSL3 or erastin treatment, DAB staining showed weak ROS accumulation only within the first invaded rice cells (Fig. 6d), which likely already proceeded to the necrotrophic stage. However, when RSL3 or erastin was applied, the DAB-positive material showed intense accumulation along fungal hyphae in all the invaded rice cells (Fig. 6d), suggesting a strong host defense response targeting the invasive hyphae. Based on these results we conclude that the induction of ferroptosis in susceptible rice plants leads to the defense response against the blast pathogen in an otherwise compatible interaction.

**Fig. 6.**
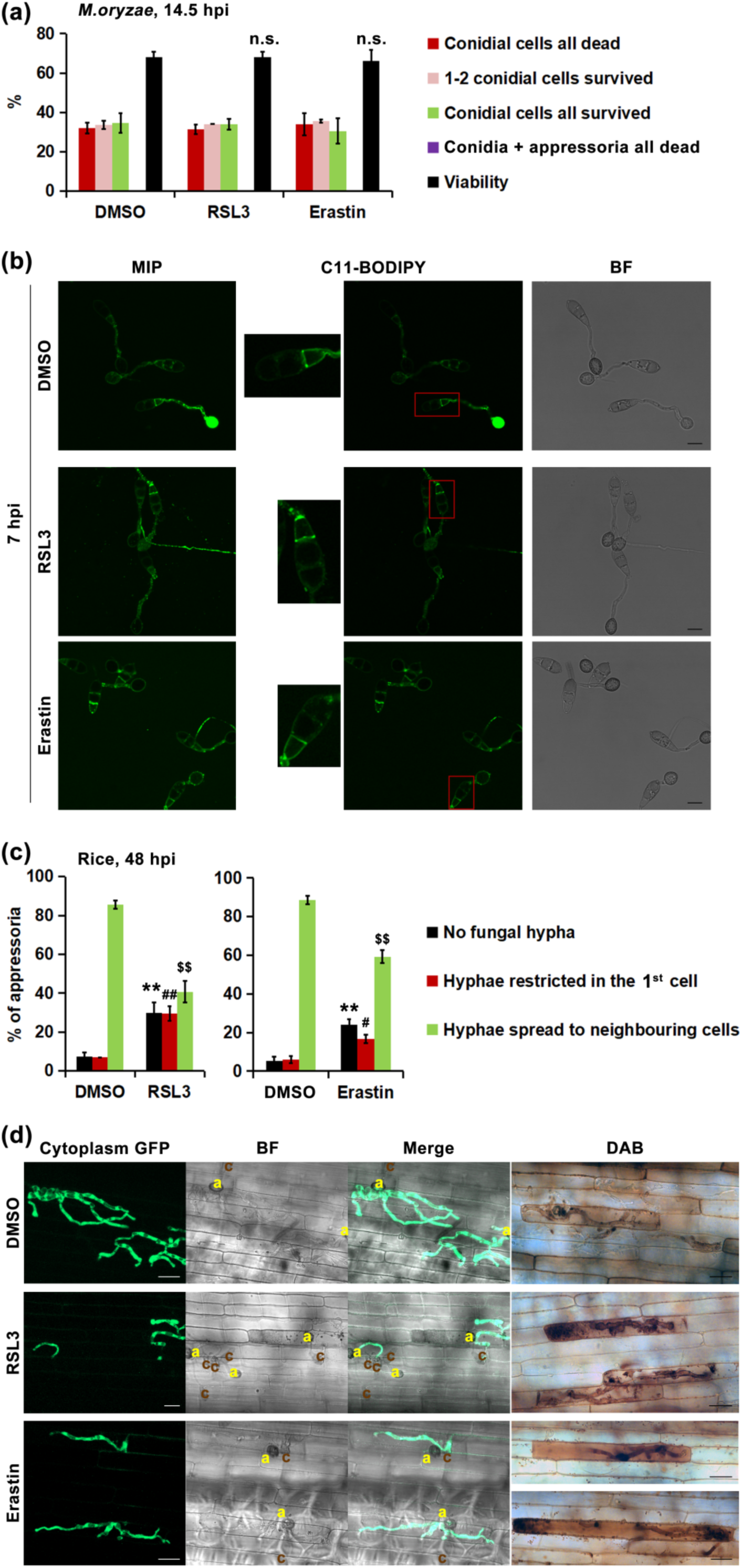
Induction of ferroptosis in susceptible rice cultivar leads to resistance against the blast fungus. (a) Viability of wild-type *M. oryzae* conidial cells treated with RSL3 or erastin at 14.5 hpi. Mean ± s.d. (3 replicates, 100 conidia per replicate) is shown; n.s., not significant versus DMSO (*t*-test). WT, wild-type. (b) Lipid oxidation, visualized by C11-BODIPY581/591 staining, in wild-type conidial cells treated with RSL3 or erastin at 7 hpi. Insets show enlarged regions-of-interest outlined in red. MIP, maximum intensity projection. C11-BODIPY, oxidized C11-BODIPY. BF, bright field. Bar = 10 **µ**m. (c) Ability of ferroptosis-inducers RSL3 and erastin to enhance rice resistance towards blast. Hyphal development of the wild-type *M. oryzae* within susceptible host (CO39) cells was quantified at 48 hpi. Each ferroptosis inducer was added at 22 hpi prior to fungal invasion, to maximize the enhancing effect on ferroptosis in the host plants (rice). Mean ± s.d. from three independent replicates (100 conidia per replicate) is shown; ** *P* < 0.01, ## *P* < 0.01 and # *P* < 0.05, and $$ *P* < 0.01 versus “no hyphae”, “hyphae restricted in the 1^st^ cell”, and “hyphae spread to neighbouring cells” of the corresponding DMSO, respectively (*t*-test). (d) Microscopic analyses of infection hyphae (*cytoplasmic GFP strain*) and ROS accumulation (DAB staining) within the rice cells at 48 hpi. BF, bright field. Merge, merge of GFP and BF images. Bar, 20 **µ**m. Data shown represents three independent experiments (biological replicates), and total n=900 conidia were used for data interpretation.

### Functional interdependency between ferroptosis and autophagy in *M. oryzae*

Autophagy has been reported to be essential for *M. oryzae* conidial cell death (Veneault-Fourrey *et al*., 2006), and has separately been shown to regulate ferroptotic cell death by modulating iron availability in mammalian cells (Gao *et al*., 2016; Hou *et al*., 2016). Therefore we set out to investigate potential links between autophagy and ferroptosis in terms of controlling conidial cell death in *M. oryzae*. *MoATG8*, a gene encoding the well-established key component of the autophagy pathway (Veneault-Fourrey *et al*., 2006; Deng *et al*., 2009), was deleted, and content of available iron was measured in the *Moatg8*Δ (hereafter referred to as *atg8*Δ) conidia. In wild-type conidia, calcein-AM fluorescence intensely accumulated in the germinating terminal cell and showed weak accumulation in the remaining two conidial cells as well as in the germ tube at 2 hpi (Fig. 7a), a time point when autophagy is robust (Fig. 7d). However, in *atg8*Δ cells, all the three conidial cells showed intense calcein-AM fluorescence (Fig. 7a), implying that *atg8*Δ conidial cells have less iron available to chelate calcein-AM. Such iron shortage was consistently observed until 24 hpi, when a small portion of *atg8*Δ conidial cells start to show the initiation of cell death (Figs S3a, 7b; n = 900). At this time point, a faint calcein-AM signal was observed in the conidium of *atg8*Δ with a maturing appressorium (Fig. S3a), which implies that irons finally accumulate to a requisite level, likely through an autophagy-independent mechanism, and can subsequently lead to death in the conidial cells. Alternatively, the faint calcein-AM signal may indicate that these *atg8*Δ conidial cells are undergoing cell death since calcein-AM is also a vital dye, and this hypothesis agrees with the finding that no calcein-AM signal was detected in dead wild-type conidial cells (Figs S3a, 7b). Consistent with calcein-AM staining, iron content in conidia at 0 hpi was much lower in the *atg8*Δ mutant (Fig. S3c). In addition, the typical membrane associated C11-BODIPY accumulation observed in wild-type conidial cells undergoing ferroptosis (Fig. 2a) disappeared upon loss of Atg8 function (Fig. S3b). Instead, punctate C11-BODIPY accumulation within *atg8*Δ conidia was observed, and the punctate fluorescence showed no response to neither the ferroptosis inducer BSO nor the inhibitor CPX (Fig. S3b). The conidium viability of *atg8*Δ moderately reduced at 24 hpi upon exogenous addition of ferric ion (FeCl_3_; *P* < 0.01, n = 900) but remained unchanged upon BSO treatment (*P* >0.05, n = 900) (Fig. 7b). This is in line with our previous finding that iron, but not BSO, attenuated CPX effect on conidial viability (Fig. 1e, g) and suggests that autophagy likely fine-tunes conidial sensitivity towards ferroptosis by modulating the availability of cellular iron. On the other hand, when CPX or Lip-1 was applied, the conidial cell death of *atg8*Δ could be further arrested, when observed at 30 hpi, reminiscent of what was seen in wild-type or the genetically complemented *atg8*Δ strain (Fig. 7c; *P* < 0.01, n = 900). We infer that iron levels in the *atg8*Δ conidia, although much lower than that in the wild-type conidia, could still support a basal albeit protracted level of ferroptosis that could be further suppressed by the iron chelator CPX or by reducing lipid peroxidation with Lip-1. Alternatively, ferroptosis may still be functional even in the absence of the appropriate autophagic activity, and the requisite iron sourced through autophagy-independent mechanisms.

**Fig. 7.**
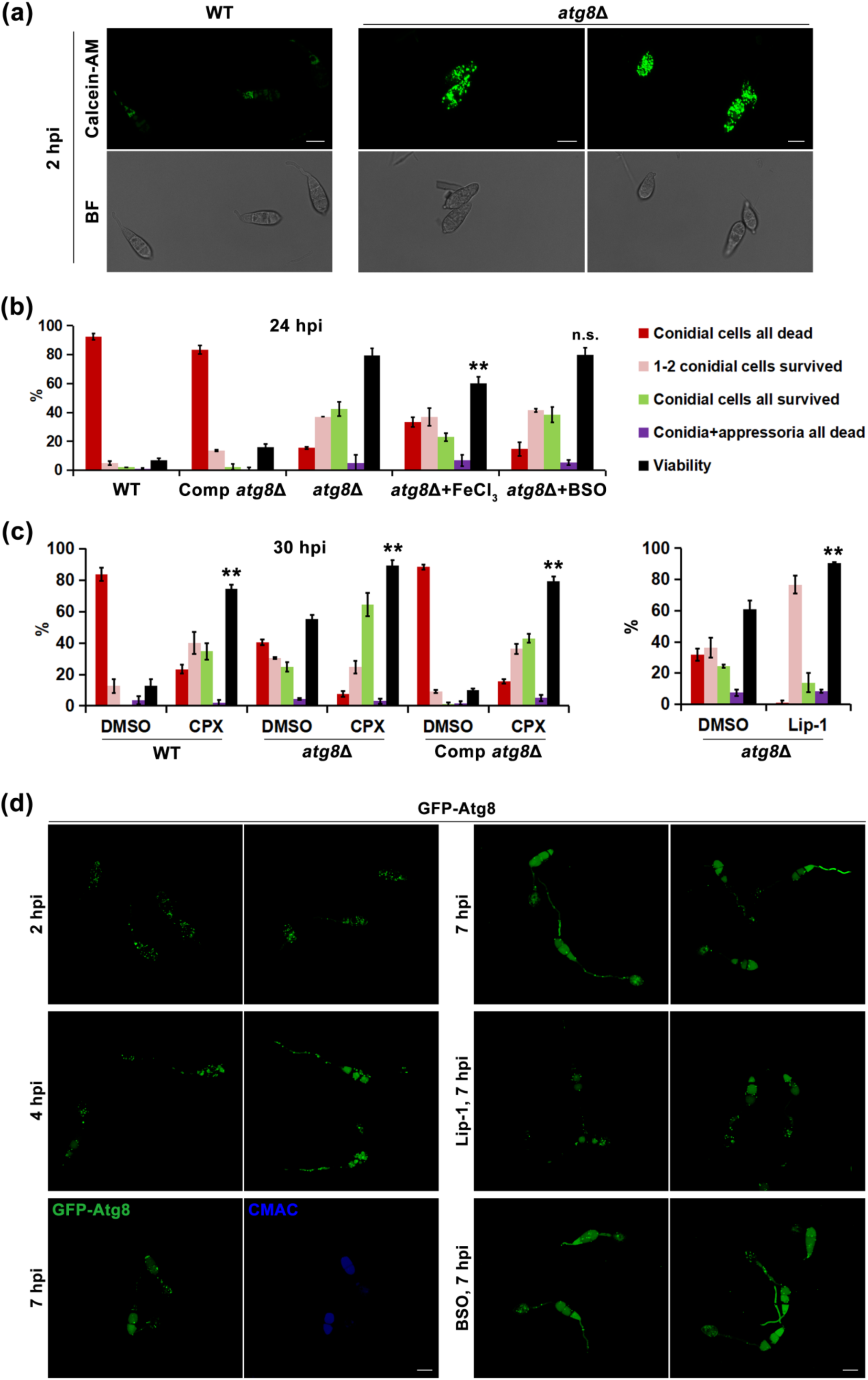
Inter-dependency between ferroptosis and autophagy in *M. oryzae*. (a) Available iron, visualized by calcein-AM staining at 2 hpi, within conidia of wild type or *atg8*Δ mutant. Calcein-AM images are shown as maximum intensity projections. BF, bright field. WT, wild-type. (b) Conidial viability assessed in the wild type, the complemented *atg8*Δ strain, or *atg8*Δ upon treatment with FeCl_3_ or BSO. Conidial viability was quantified at 24 hpi, and is plotted as mean ± s.d. (3 replicates, 100 conidia for each replicate); ** *P* < 0.01 and n.s. (not significant) versus *atg8*Δ (*t*-test). WT, wild-type. (c) Ability of CPX and Lip-1 to increase conidial viability of wild-type, *atg8*Δ, and the complemented *atg8*Δ strain at 30 hpi. Mean ± s.d., 3 replicates (100 conidia per replicate); ** *P* < 0.01 versus corresponding DMSO control. WT, wild-type. (d) Subcellular localization of GFP-Atg8 in the presence or absence of Lip-1 or BSO during conidial development (2, 4, 7 hpi). CMAC staining marks the fungal vacuoles. All data are representative of two to three independent biological replicates, and total n=900 conidia were used for viability interpretation. Bar = 10 **µ**m for all the images.

We further investigated the relationship between autophagy and ferroptosis by examining autophagy dynamics (visualized using GFP-Atg8) in response to treatment with ferroptosis inducers or inhibitors at multiple time points. We could see that autophagy was induced in the wild-type conidia as early as 2 hpi on inductive surface, and proceeded as GFP-Atg8 accumulated in the vacuoles at 4-7 hpi (Fig. 7d). At 7 hpi, most of GFP-Atg8 signals were vacuolar, indicating that autophagy completed in the wild-type conidia (Fig. 7d), when ferroptosis initiated (Fig. 3c-f). However, no significant change in autophagy processes (and by implication function/activity) was observed regardless of enhancing or suppressing the ferroptosis (Figs 7d, S3d). Taken together, we conclude that autophagy regulates conidial ferroptosis by modulating the availability of intracellular iron, but neither the enhancement nor the suppression of ferroptosis seem to affect autophagy *per se* during appressorial development in *M. oryzae*.

## Discussion

Ferroptosis is a regulated cell death pathway implicated in several disease contexts in mammals (Stockwell *et al*., 2017), heat stress in Arabidopsis (Distéfano *et al*., 2017), and in specific effector-triggered immunity in rice (Dangol *et al*., 2019). In this article, we have shown that ferroptosis is also involved in developmental cell death in *M. oryzae* conidia and that such ferroptotic cell death is necessary for proper rice blast infection. A correlation between ferroptotic conidial death and appressorium development was observed (Figs 1c, 2a, 4b, S1b,d), implying that conidial ferroptosis may modulate the proper formation and maturation of the specialized infection structure to regulate the blast disease process. In addition, we also provided some important insight into the underlying mechanism(s) in the control of conidial ferroptosis and its contribution to *M. oryzae* pathogenicity.

We showed that *M. oryzae* employs the superoxide-producing Nox enzymes as important sources of ferroptosis-driving lipid peroxidation (Figs 4, S1h), just like what was reported in tumor cells (Dixon *et al*., 2012). However, an important dissimilarity that needs to be emphasized is that the biological context for ferroptosis found in tumor cells is different compared to that observed in *M. oryzae*. The tumor cells, wherein major ferroptosis mechanisms were identified, are mammalian cells bearing an activated Ras mutation, and successful ferroptosis/cell death therein requires inducers like erastin, RSL3, or BSO (Yang & Stockwell, 2008; Yang *et al*., 2014). However, conidial cells, in which ferroptosis was identified in this study, represent unperturbed wild-type *M. oryzae* disease propagules that undergo cell death as well as the oxidative processes naturally for infection-related morphogenesis and development. These differences likely explain why lipid peroxidation was observed in *M. oryzae* germ tubes and maturing appressoria (Figs 2a, 3a, 4f, 6b), which are viable and metabolically active. In addition, these differences may also explain why BSO indeed sensitized conidial cells to cell death, highly likely through GPX inhibition, but the mammalian GPX4-specific inhibitor RSL3 showed no effect on ferroptosis in *M. oryzae* (Figs 1g, 2a, 6a,b), probably because the binding site in the target protein for the small molecule RSL3 is not conserved in *M. oryzae*. Recently, two studies described a GPX4-independent mechanism that dominantly controls ferroptosis in tumor cells that are insensitive to GPX4 inhibition (Bersuker *et al*., 2019; Doll *et al*., 2019). It will be interesting to test whether such GPX4-independent pathway functions in *M. oryzae* too.

In a seminal study, ferroptosis was reported to be involved in the HR response during an incompatible interaction with rice blast (Dangol *et al*., 2019). However, this study did not envisage or account for ferroptosis in *M. oryzae* when designing the experiments, and all the treatments were applied to both *M. oryzae* and rice cells simultaneously during the incompatible interaction. However, taking our findings into consideration, we do believe that in addition to such contribution to the HR response during the incompatible interaction, ferroptotic cell death plays a key role in proper pathogenic development *per se* in the blast fungus too. Our results suggest that ferroptosis is suppressed or is at a minimal basal level in the blast-susceptible rice plants, therefore susceptible rice is colonized successfully by *M. oryzae*. However, when induced by specific chemicals like RSL3 or erastin, ferroptosis and the associated immune defense is greatly enhanced and robustly attacks the invasive hyphae to restrict the development and spread of rice blast in an otherwise susceptible cultivar. Overall, we think that ferroptosis is a conserved cell death mechanism employed by both the partners involved in the blast pathosystem albeit for entirely different purposes or outcomes. Autophagy was reported to be essential for conidial cell death, which is a prerequisite for appressorium functioning and host infection (Veneault-Fourrey *et al*., 2006). Our results showed that loss of autophagy (the *atg8*Δ mutant) caused increased accumulation of lipid peroxides on vesicular membranes within the *atg8*Δ conidia, amidst a cellular environment with limited availability of iron (ferrous and ferric ions) during pathogenesis (Fig. S3a,b,c). Therefore, we infer that, in addition to directly controlling the conidial cell death, autophagy indirectly regulates ferroptosis by modulating the iron availability. This is further supported by our finding that addition of FeCl_3_ partially bypasses autophagy function and induces/promotes conidial cell death (Fig. 7b). The extremely low levels of intracellular iron in the *atg8*Δ conidia, probably achieved through autophagy-independent mechanism, may support a basal level of ferroptosis (conidial death), which could be further inhibited by either chelating iron with CPX, or by reducing lipid peroxidation with Lip-1 (Figs 7a-c, S3a). Further experiments focusing on iron homeostasis will help elucidate the relationship and codependence between autophagy and ferroptosis.

## Acknowledgements

We thank the Fungal Patho-biology Group (TLL) for helpful discussions and suggestions. We thank Kou Yanjun for the *GFP-ATG8* strain. This study was funded by grants from the Temasek Life Sciences Laboratory (Singapore) to N.I.N; and the National Natural Science Foundation of China (no. 31970139) to D.Y.Z.

## Author Contributions

S.Q, Y.Z.D. and N.I.N. designed the study. S.Q, M.L and Y.F performed the experiments. S.Q, Y.Z.D and N.I.N analyzed the data; and co-wrote and revised the manuscript.

## Supporting Information

**Fig. S1** Ferroptosis contributes to cell death during *M. oryzae* pathogenesis.

**Fig. S2** Ferroptotic fungal cell death is required for *M. oryzae* pathogenicity.

**Fig. S3** Crosstalk between ferroptosis and autophagy in the conidium of rice blast fungus.

**Table S1** Oligonucleotide primers used for plasmid constructs.

