## Supplementary Figures for "Ferroptosis contributes to developmental cell death in rice blast"

**Supplementary data (Shen et al.)**

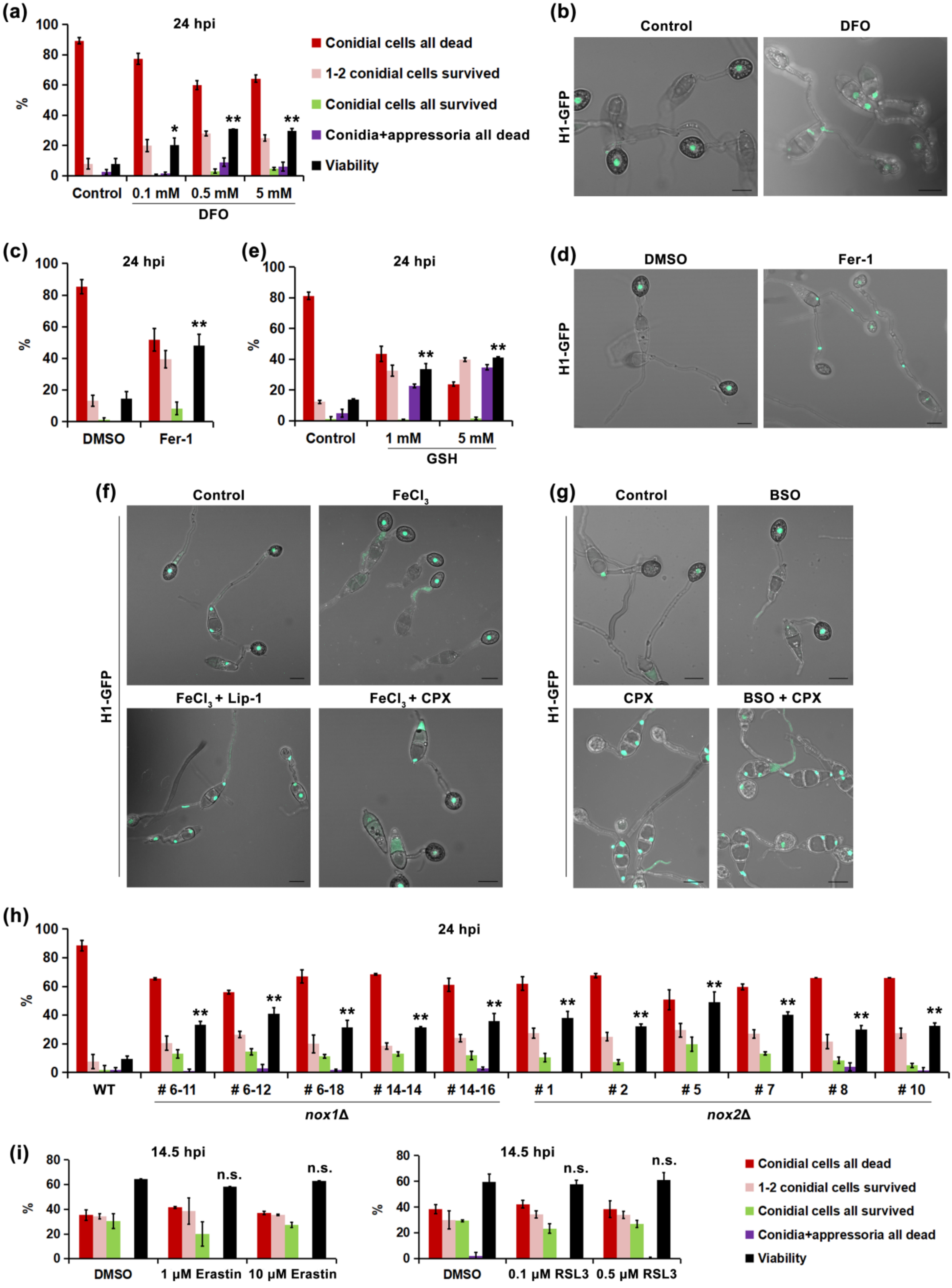

**Fig. S1** Ferroptosis contributes to cell death during *M. oryzae* pathogenesis. (a) Ability of iron chelator DFO to prevent conidial death. Conidial viability (visualized using a *HistoneH1-GFP* strain) was measured at 24 hpi. Mean ± s.d. from three independent experiments (100 conidia for each experiment) is shown; * $P < 0.05$ and ** $P < 0.01$versus control (*t*-test). (b) Epifluorescence microscopy to visualize conidial viability (*H1-GFP* strain) at 24 hpi, in the presence of DFO. (c) Ability of Fer-1 to prevent conidial death, measured at 24 hpi using the *H1-GFP* strain. Mean ± s.d., 3 replicates, 100 conidia per replicate; ** $P$ <0.01versus DMSO (*t*-test). (d) Visualization of conidial viability of *H1-GFP* strain ± Fer-1 treatment at 24 hpi using epifluorescence microscopy. (e) Viability of *M. oryzae* cells after GSH treatment, measured at 24 hpi using the *H1-GFP* strain. Mean ± s.d., 3 replicates, 100 conidia for each replicate; ** $P < 0.01$versus control (*t*-test). (f) Visualization of conidial viability, at 24 hpi using epifluorescence microscopy, of $FeCl_3$ treated *H1-GFP* strain in the presence or absence of Lip-1 or CPX. (g) Confocal microscopy to visualize conidial viability of BSO treated *H1-GFP* strain in the presence or absence of CPX at 24 hpi. (h) Conidium viability of wild-type and *nox*Δ mutants at 24 hpi. Mean ± s.d., 3 replicates (each has 100 conidia); ** $P < 0.01$ versus WT (*t*-test). WT, wild-type. (i) Conidium viability of wild-type *M. oryzae* conidial cells treated with RSL3 or erastin at 14.5 hpi. Mean ± s.d. from three replicates (n=100) is shown;  n.s., not significant versus corresponding DMSO (*t*-test). All the H1-GFP images are shown as maximum intensity projections. The experiments were repeated at least twice, with n=900 or n=600 conidia in total for data interpretation. Bars equal 10 $\mu$m.

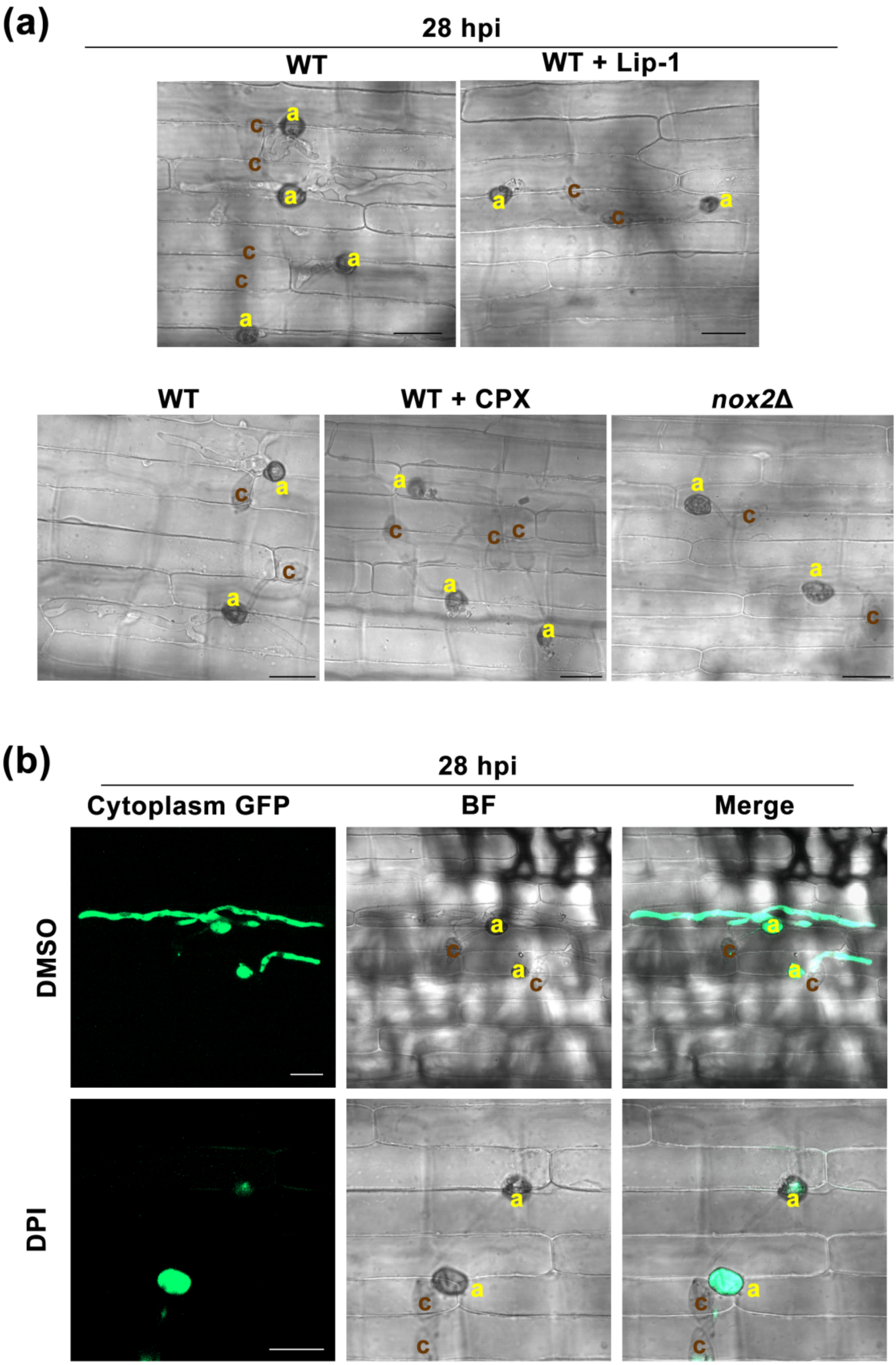

(a)
28 hpi
WT
WT + Lip-1
WT
WT + CPX
nox2Δ
(b)
28 hpi
Cytoplasm GFP
BF
Merge
DMSO
DPI

**Fig. S2** Ferroptotic fungal cell death is required for *M. oryzae* pathogenicity. (a) Ability of wild-type or *nox2*Δ ± Lip-1or CPX treatment to develop infection hyphae within the susceptible host plants (CO39) at 28 hpi. C, conidia; A, appressorium. WT, wild-type. (b) Microscopy observation of infectious development within rice cells at 28 hpi using the blast strain expressing the cytosolic GFP. BF, bright field. Merge, merge of GFP and BF images. C, conidia; A, appressorium. All the ferroptosis inhibitors were washed away at 22 hpi to restrict the chemical effect on conidial death. Scale bar equals 20 micron. Experiments reported here were performed as three independent biological replicates.

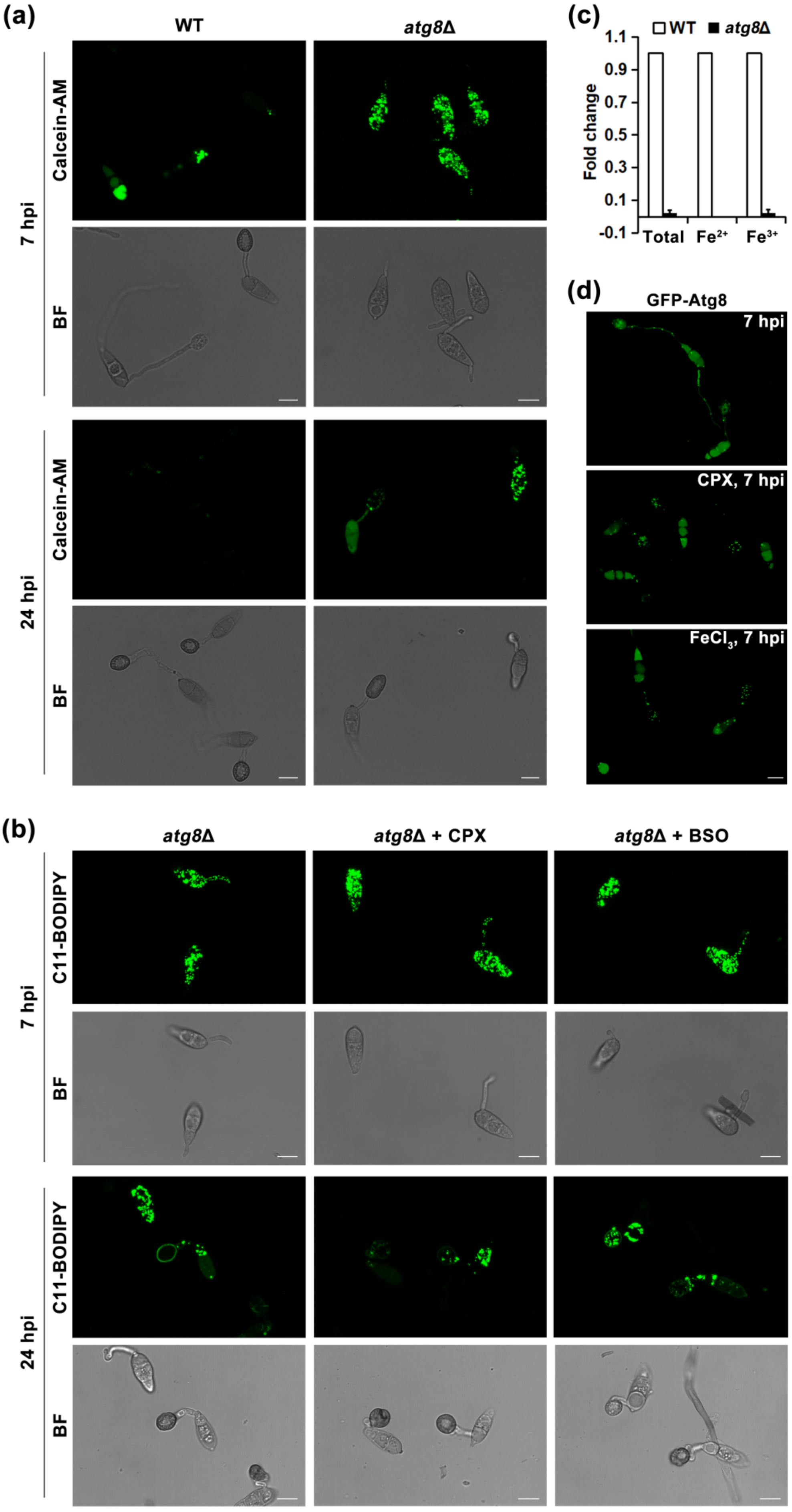
(a)
WT
atg8Δ
7 hpi
Calcein-AM
BF
24 hpi
Calcein-AM
BF
(c)
WT
atg8Δ
Fold change
1.1
0.9
0.7
0.5
0.3
0.1
-0.1
Total
Fe2+
Fe3+
(d)
GFP-Atg8
7 hpi
CPX, 7 hpi
FeCl3, 7 hpi
(b)
atg8Δ
atg8Δ + CPX
atg8Δ + BSO
7 hpi
C11-BODIPY
BF
24 hpi
C11-BODIPY
BF

**Fig. S3** Crosstalk between ferroptosis and autophagy in the conidium of rice blast fungus. (a) Cellular iron levels, visualized by calcein-AM staining at 7 and 24 hpi, in wild-type and *atg8*Δ. BF, bright field. WT, wild-type. (b) C11-BODIPY581/591 staining showing lipid peroxidation in *atg8*Δ conidial cells upon treatment with CPX or BSO at 7 and 24 hpi. BF, bright field. C11-BODIPY, oxidized BODIPY581/591. (c) Measurement of iron content (total Fe, $Fe^{2+}$, $Fe^{3+}$) in wild type and *atg8*Δ conidia (0 hpi). Total iron content is presented as fold change relative to that in the wild type strain. Data are mean ± s.d. derived from two independent biological repeats, each of which contains two technical repeats, with $3\times10^5$ conidia in each measurement. WT, wild-type. (d) GFP-ATG8 subcellular localization in the absence or presence of CPX or $FeCl_3$ at 7 hpi. Except for GFP-Atg8, all the images are shown as maximum intensity projections. Scale bar marks 10 micron. Data shown were derived from 2 independent experiments.

**Table S1** Oligonucleotide primers used for plasmid constructs

| Gene (Locus) | Description | Application | Primer sequence (Enzyme site underlined) |
|---|---|---|---|
| *Histone H1* (*MGG_12797*) | GFP-tagging (C-terminal) | GFP without start codon | 5'-CATCGGTACCGTGAGCAAGGGCGAGGAGCTGT-3'<br>5'-CGCGGATCCTTACTTGTACAGCTCGTCCATGC-3' |
| | | 5' homologous arm (ORF without stop codon) | 5'-CGTCTCGAGCCTCCCAAGAAGGAAACC-3'<br>5'-GTCGAATTCTGCGGCGGGTGCCTCGGC-3' |
| | | 3' homologous arm | 5'-CCGCTGCAGTAAAGGGACGCTGACGAACTT-3'<br>5'-CATAAGCTTCTTTCTTTGACGGGAAAGGGA-3' |
| *Histone H3* (*MGG_01159*) | Cytoplasm GFP under *Histone H3* promoter | Histone H3 | 5'-TGTGAATTCGTGGGGGACGACCTTACCT-3'<br>5'-CACGGTACCCATGGTGATTGATTTGTGATTGATG-3' |
| | | GFP without start codon | 5'-CATCGGTACCGTGAGCAAGGGCGAGGAGCTGT-3'<br>5'-CGCGGATCCTTACTTGTACAGCTCGTCCATGC-3' |
| *NOX1* (*MGG_00750*) | Deletion construct | 5' and 3' homologous arms | 5'-GCCGGTACCCAAGTCCATGTGGCAGTAGTTC-3'<br>5'-GCCGGATCCGATACAAGCAAAACAAGCGACC-3'<br>5'-GCCCTGCAGAGAAAAGCTCGGGAAGGTGG-3'<br>5'-GCCAAGCTTCATCAACGGTTTATGTACGGATTG-3' |
| *NOX2* (*MGG_06559*) | Deletion construct | 5' and 3' homologous arms | 5'-GCCGGTACCCGGTGATACGGTCTCCATAAGTCG-3'<br>5'-GCCGGATCCCAGCTTACAACGGAGACGTTCG-3'<br>5'-GCCCTGCAGGTAGTTATGGGCAGGTCT-3'<br>5'-GCCAAGCTTCTACCAGAGGCAGTACAAGG-3' |
